# Pooled endogenous protein tagging and recruitment for scalable discovery of effectors for induced proximity therapeutics

**DOI:** 10.1101/2023.07.13.548759

**Authors:** Yevgeniy V. Serebrenik, Deepak Mani, Timothé Maujean, George M. Burslem, Ophir Shalem

## Abstract

The field of induced proximity therapeutics is in its ascendancy but is limited by a lack of scalable tools to systematically explore effector-target protein pairs in an unbiased manner. Here, we combined Scalable POoled Targeting with a LIgandable Tag at Endogenous Sites (SPOTLITES) for the high-throughput tagging of endogenous proteins, with generic small molecule-based protein recruitment to screen for novel proximity-based effectors. We apply this methodology in two orthogonal screens for targeted protein degradation: the first using fluorescence to monitor target protein levels directly, and the second using a cellular growth phenotype that depends on the degradation of an essential protein. Our screens revealed a multitude of potential new effector proteins for degradation and converged on members of the CTLH complex which we demonstrate potently induce degradation. Altogether, we introduce a platform for pooled induction of endogenous protein-protein interactions that can be used to expand our toolset of effector proteins for targeted protein degradation and other forms of induced proximity.

## Introduction

Chemically induced proximity is a rapidly advancing field aimed at expanding the druggable genome beyond inhibition of enzymatic targets^1^. The most prominent and advanced type of induced proximity modality is targeted protein degradation (TPD), involving the depletion of a target protein through proximity to an effector protein with degradative function. Typical effectors include E3 ligases that trigger target ubiquitination and subsequent degradation via the ubiquitin proteasome system (UPS)^2^. Proximity is induced by small molecules like proteolysis targeting chimeras (PROTACs) or molecular glues that bind between the effector and target protein^3^. TPD-based therapies offer numerous advantages over traditional pharmacology including the complete abrogation of pathogenic proteins and the expansion of the druggable protein space by leveraging any viable binding site on a protein. Proximity-based degraders are not limited to action through the UPS, and initial reports have shown the autophagy lysosome pathway (ALP) as another potential option for inducing protein degradation^4,5^. Effectors with other functions, such as protein stabilizers, kinases, or phosphatases, are also being developed for appropriate therapeutic applications^6–9^.

One major limiting factor in the fast development of this new type of therapeutic modality is that even in TPD, the number of available effector proteins is extremely limited. Over 90% of developed PROTACs recruit CRBN or VHL, while the rest use one of approximately 10 other E3 ligases^10,11^ For other modalities, the list is even more limited and is largely determined by the availability of ligands rather than the optimal nature of the recruited protein. This leaves hundreds of unexplored potential effector proteins including over 700 and 200 proteins that belong to the UPS and ALP, respectively, and hundreds of others that belong to additional branches of the proteostasis network^12,13^. Exploring this large effector space will be essential for the successful implementation of proximity-based therapeutics in the clinic across multiple disease areas. A major advantage of orthogonal effectors, particularly ones that are growth-essential, would be to overcome common resistance mechanisms to TPD cancer therapies including effector downregulation, binding mutations, or mutations in other UPS components that impair degradation^14–16^. In cases where a target is inefficiently degraded by the current suite of effectors, novel effectors with more compatible structures, localizations, or expression patterns are necessary^17^. In addition, effectors with tissue-specific expression represent the holy grail in TPD since they avoid systemic off-target effects^18^. Finally, therapeutic approaches beyond TPD enable diverse actions such as protein stabilization, disaggregation, or specific post-translational modifications, thus enabling new swaths of biology to be modulated. A catalog of proteins with different mechanisms, expression patterns, and effector functions would thus be invaluable for the expansion of proximity-based therapeutics into new modalities.

The utilization of new effectors has been slow, relying on existing chemistry for recruitment that has been discovered through serendipity^19,20^, rational design^21,22^, or focused chemical profiling^23–26^. More systematic approaches for the characterization of protein function by induced proximity are starting to be realized using ectopically expressed protein fragments^27^ or open reading frames^28^ fused to generic recruitment domains to assess their transcriptional or protein degradation activity. These approaches have been successful in the elucidation of protein function at scale, yet previous attempts to use effector overexpression for TPD discovery has yet to produce efficient small molecule degraders, potentially explained by large differences in expression levels or stoichiometry with endogenous cofactors^29^.

In this study, we present a novel platform for the systematic and scalable profiling of effector function by endogenous protein recruitment. We utilized our recently developed method, Scalable POoled Targeting with a LIgandable Tag at Endogenous Sites (SPOTLITES), to generate a cell library with hundreds of potential effector proteins that are tagged with HaloTag at multiple locations^30^. We then used optimized heterobifunctional small molecules for endogenous protein recruitment to systematically assess the degradation potential of effector proteins in two parallel approaches: fluorescence-based monitoring of target protein expression and a growth-based approach capitalizing on the cellular toxicity induced by BRD4 degradation^31–33^. These experiments resulted in the discovery of multiple effectors for potential use in TPD. Excitingly, both approaches converged on proteins within the CTLH E3 ligase complex. We further show that recruitment of several proteins within this complex results in rapid and efficient degradation of targeted substrates, and that such degradation depends on other proteins within the complex. These data, together with studies that show the importance of CTLH complex expression in cancer progression^34^, make CTLH complex members ideal new effectors for TPD. More importantly, we present a novel platform for rapid, highly scalable, endogenous protein recruitment which can be applied to profile effectors for any type of protein modification in various cell lines, facilitating the accelerated development of proximity-based therapeutics.

## Results

### Optimizing a generic approach for inducing ectopic protein-protein interactions

To survey effector function at a proteome scale, we set to design a flexible approach to generically induce protein-protein interactions between a target of interest and many endogenously tagged potential effector proteins simultaneously. This approach is based on our recent method, Scalable POoled Targeting with a LIgandable Tag at Endogenous Sites (SPOTLITES)^30^, in which we construct a complex cell pool where endogenous proteins are tagged with HaloTag (HT), a generic ligand-binding domain^35^. Each cell in the pool has a single protein site tagged with HT that can be recruited to a target with an orthogonal ligand binding domain by treatment with a generic heterobifunctional molecule (Fig. 1A). We first tested the degradation of cytonuclear targets using a model protein, mClover3 fused to FKBP^F36V^ (mClover3-FKBP)^36^, that would allow for a direct readout of degradation by fluorescence after treatment with ligands that bind both HT and FKBP^36^. We then used flow cytometry-based assays to optimize ligand and cell line conditions on a clonal HAP1 cell line stably expressing mClover3-FKBP in both the nucleus and cytosol (Fig. 1B).

**Figure 1.**
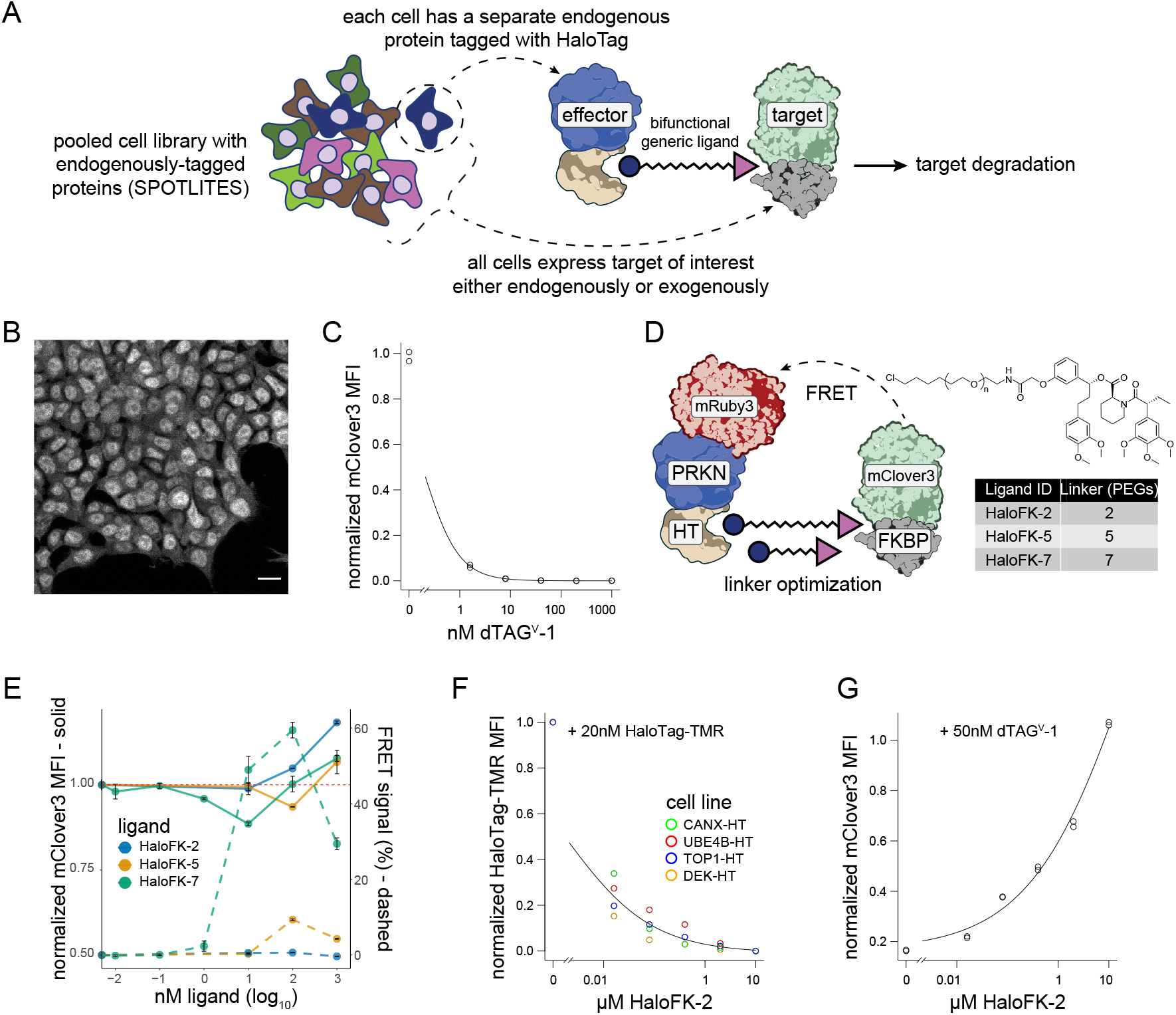
Optimizing a generic approach for inducing ectopic protein-protein interactions. A) Diagram depicting strategy for pooled induction of protein-protein interactions to find endogenous effectors for targeted protein degradation using a SPOTLITES library. B) Confocal imaging of mClover3-FKBP expression in a clonal HAP1 cell line. Scale bar represents 10 μm. C) Degradation of mClover3-FKBP after 32 hour treatment with dTAG^V^-1 as measured by flow cytometry. Two replicates are shown for each concentration. D) Overview of optimization approach for HaloTag-FKBP heterobifunctional ligands with various linker lengths, using mRuby3-PRKN-HaloTag as a model effector. E) Effect of HaloTag/FKBP heterobifunctional ligands on mClover3-FKBP levels (left axis) or FRET with mRuby3-PRKN-HaloTag (right axis). F) Competition of HaloFK-2 with HaloTag-TMR for binding to HaloTag in four clonal, endogenously-tagged HaloTag cell lines. G) Competition of HaloFK-2 with dTAG^V^-1 for binding to FKBP^F36V^ as measured by degradation of mClover3-FKBP. Two replicates are shown for each concentration.

We assessed the ability to degrade this target with endogenous degradation machinery by treatment with dTAG^V^-1, a potent FKBP-based PROTAC that recruits the VHL E3 ligase^37^. Even at low nanomolar concentrations of dTAG^V^-1, mClover3-FKBP was almost completely depleted after 24 hours, validating its viability as a substrate for targeted protein degradation (Fig. 1C). To determine the optimal HT/FKBP bifunctional ligand linker length as well as treatment concentration, we introduced a model effector, the E3 ligase PRKN, that was previously shown to function exogenously^29^. PRKN was fused to HT for binding the generic ligand, and to mRuby3 to assess ternary complex formation between the effector, ligand, and target by fluorescence resonance energy transfer (FRET) (Fig. 1D). Fortuitously, mRuby3-PRKN-HT induced only minor degradation of mClover3-FKBP, allowing us to simultaneously measure degradation and ternary complex formation efficiency in the same system in response to heterobifunctional ligands of different lengths (Fig. 1D). The 7-PEG bifunctional ligand, HaloFK-7 (also known as PhosTAC7^38^), demonstrated strong FRET signal at the same concentrations that induced degradation (Fig. 1E). In contrast, the shorter 5-PEG ligand, HaloFK-5, induced both a weaker FRET signal and less degradation (Fig. 1E). The shortest ligand, HaloFK-2, induced neither FRET nor detectable degradation, leading only to the stabilization of the target protein (Fig. 1E), suggesting that longer linkers are needed for ternary complex formation and subsequent induced degradation.

To confirm that HaloFK-2 was still able to bind both intended targets, we used orthogonal competition assays. Stable cell lines expressing HT-tagged proteins labeled with HT-TMR were pretreated with HaloFK-2 at various concentrations, leading to reduced TMR signal (Fig. 1F). Similarly, dTAG^V^-1-mediated depletion of mClover3-FKBP was mitigated by pretreatment with HaloFK-2 (Fig. 1G). Together, these results indicate that while HaloFK-2 is able to bind both HT and FKBP separately, it is not able to bind both simultaneously, most likely due to its short linker length. We thus decided to employ HaloFK-7 to study induced proximity of targets with endogenously tagged proteins and HaloFK-2 as a control to account for indirect effects of binding both the effectors and targets.

### Comparison of two orthogonal effector screens highlight the CTLH complex for its potential to degrade recruited substrates

Having optimized conditions for potent and generic induced degradation, we set out to perform screens using a large number of endogenously tagged potential effectors. We chose to initially tag components of the proteostasis machinery, including 569 genes involved in the ubiquitin proteasome system (UPS), 236 in the autophagy lysosome pathway (ALP), and 148 in the chaperone network (Fig. 2A). We constructed a SPOTLITES cell pool using an sgRNA library for endogenous tagging^30^ (www.pooledtagging.org). Much like for CRISPR screens^39^, the sgRNA library is transduced at low multiplicity of infection followed by antibiotic selection. Subsequently, generic reagents for tagging by homology-independent intron targeting^40^ are delivered, including the DNA tag donor encoding HaloTag flanked by splicing sequences, an sgRNA-expressing plasmid to linearize the donor, and a Cas9-expressing plasmid for DNA cutting. Cells with successful tagging are enriched by fluorescence-activated cell sorting using a fluorescent HT ligand, after which they can be assayed and easily analyzed by spacer amplicon sequencing (Fig. 2B).

**Figure 2.**
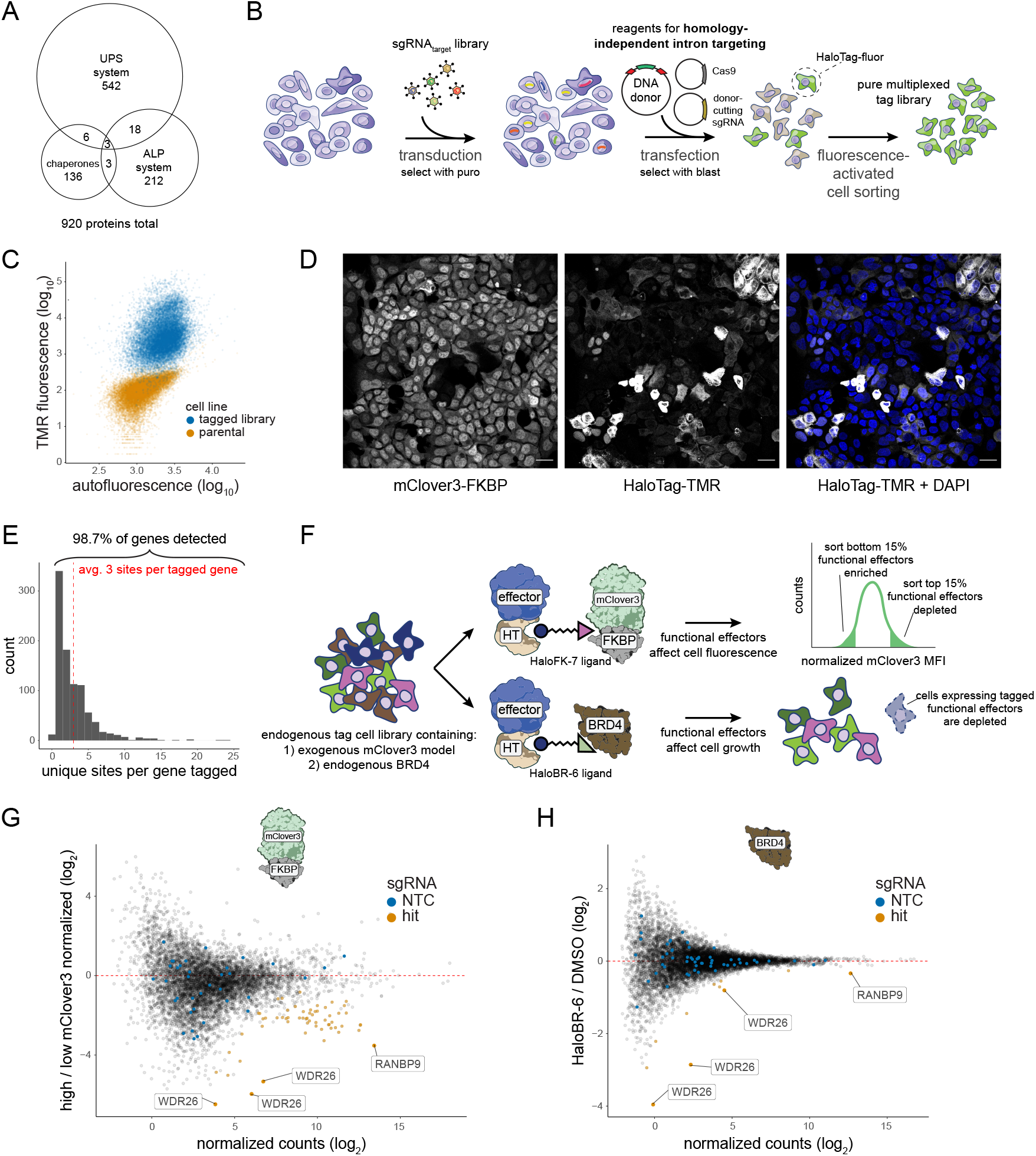
Comparison of two orthogonal effector screens highlight the CTLH complex for its potential to degrade recruited substrates. A) Euler diagram depicting the targeted effector proteins and their categorizations based on GO:CC. B) Strategy for generating a SPOTLITES library of endogenously-tagged effector proteins. C) Flow cytometry-based visualization of the pooled tag library treated with HaloTag-TMR in comparison to the parental cell line. D) Confocal imaging of the pooled tag library treated with HaloTag-TMR to visualize individual tagged proteins. Scale bars represent 20 μm. E) Histogram of the number of different fusion sites tagged for each targeted protein in the pooled library. F) Strategy for performing two orthogonal screens for functional effectors of degradation by targeting either mClover3-FKBP (top path) or endogenous BRD4 (bottom path). Degraders of mClover3-FKBP are enriched by FACS, while degraders of BRD4 are depleted during cell growth. G-H) Abundance versus fold change plots for tagging sgRNAs in the G) fluorescence- or H) growth-based screen. In G, the ratio of sgRNAs in bright versus dim mClover3 populations is compared between cells treated with HaloFK-7 or HaloFK-2. All sgRNAs representing non-targeting controls (NTCs) are colored in blue, screen hits are orange, and select hit sgRNAs are labeled with the name of the tagged protein.

Using this approach, we generated a diverse cell library in haploid HAP1 cells (Fig. S1A) of highly-expressed endogenous proteins as measured by flow cytometry with a fluorescent HaloTag label (Fig. 2C). Confocal imaging revealed cells with many different localization patterns indicative of the variety of proteins tagged (Fig. 2D). To analyze library composition in detail, we performed deep sequencing of spacers. We were able to detect almost all targeted proteins (Fig. 2E) based on an inclusive sgRNA count threshold set by background reads (Fig. S1B). As proteins were targeted with multiple sgRNAs, each protein on average was represented by 3 different tag fusion sites (Fig. 2E), allowing us to test effectors at multiple orientations to the target and obtain information on productive ternary complex geometry simultaneously with effector identity.

We used our cell library of tagged effector proteins to perform two parallel screens against different targets using two distinct heterobifunctional molecules (Fig. 2F). One target was the exogenous mClover3-FKBP which was expressed in each tagged cell (Fig. 2D). This target allowed us to screen directly on protein levels using the HaloFK-7 ligand. We also targeted endogenously-expressed BRD4 with a different ligand that we synthesized to bind both HaloTag and BRD4 directly, HaloBR-6 (Fig. S1C). BRD4 degradation has a notable anti-proliferative phenotype^31–33^, which we confirmed in our cell line using an established CRBN-based BRD4 degrader^41^ in comparison to a BRD4 inhibitor^42^ (Fig. S1D). Thus, in the BRD4 screen, we used cell depletion as an indirect measure of target degradation (Fig. 2F).

To screen for degraders of mClover3-FKBP, we treated the library with either the HaloFK-7 ligand or the negative control, HaloFK-2, for 22 hours and sorted cells based on mClover3 intensity (Fig. S1E), followed by genomic DNA extraction and sequencing. As tagging of proteins within the proteostasis network has effects on target levels independent of recruitment (Fig. S1F-G), we compared the ratios of sgRNAs in the high and low mClover3 populations between HaloFK-7 and HaloFK-2 (Fig. S1H), revealing tags that affect mClover3 levels only in a recruitment-dependent manner (Fig. 2G). Similarly, to screen for degraders of endogenous BRD4, the cell library was treated with the HaloBR-6 ligand or vehicle control for 72 hours. sgRNAs depleted in treated versus control samples clearly revealed specific effectors of degradation (Fig. 2H). Notably, many of the top hits were shared between the two screens, suggesting that the discovered effectors of degradation are capable of targeting a variety of ectopic substrates. With these two orthogonal screens, we were able to discover multiple classes of endogenous degradation effectors, including E3 ligase substrate adaptors, E3 ligase complex scaffolds, and proteins involved in autophagy. This establishes our platform as a robust method for systematic study of induced proximity-based effectors expressed at physiological levels.

### Members of the CTLH complex can function as flexible effectors for TPD

We further investigated two strong hits from both screens, WDR26 and RANBP9, with known scaffolding roles within the CTLH E3 ligase complex, with WDR26 also acting as a potential substrate receptor^43–46^. For each of these proteins, we generated clonal endogenous tag lines in HAP1 cells using the enriched sgRNAs from the screens. After treatment with HaloFK-7 for 24 hours, both effectors substantially reduced levels of mClover3-FKBP with slightly bimodal single cell profiles (Fig. 3A,B). While most WDR26-tagged cells exhibited complete degradation, rivaling that with dTAG^V^-1 (Fig. 3A), the majority of RANBP9-tagged cells showed ∼40% degradation, with a smaller population exhibiting near-complete degradation (Fig. 3B). Target degradation via endogenously-tagged WDR26 was also potent in HEK293 cells, demonstrating that this effector is not cell line-specific (Fig. S2A). Dose response experiments confirmed that the concentration used for screening was the degradation maximum, consistent with our initial ligand optimization experiments (Fig. 3C). To examine degradation kinetics, time course experiments revealed that while WDR26 induced over 90% target degradation in under 24 hours, RANBP9 exhibited slower kinetics, with a steady rate of mClover3-FKBP depletion as far out as 48 hours after treatment (Fig. 3D).

**Figure 3.**
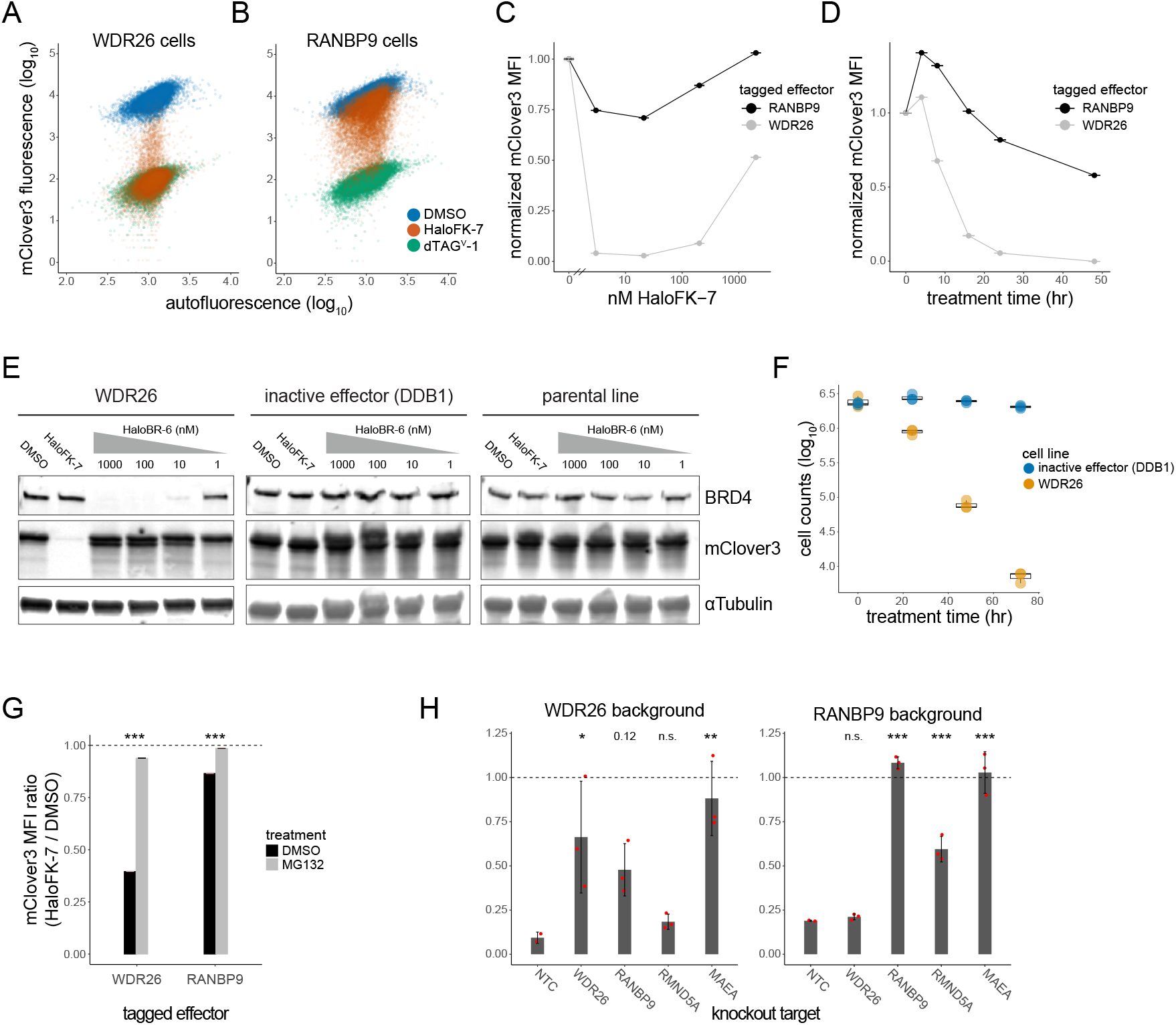
Members of the CTLH complex can function as flexible effectors for TPD. A-B) Flow cytometry plots of A) WDR26- or B) RANBP9-tagged HAP1 cells treated with the indicated drugs for 72 hours. C-D) mClover3-FKBP mean fluorescence intensity (MFI) in tagged cells treated with HaloFK-7 for C) 24 hours at indicated concentrations, or at D) 20 nM HaloFK-7 for the indicated amounts of time. E) Immunoblot representing 24 hour degradation of BRD4 or mClover3-FKBP with indicated concentrations of HaloBR-6 or 20 nM HaloBR-7, respectively, in cells with tagged WDR26, DDB1, or in untagged cells (parental). F) Counts of cells with tagged WDR26 or DDB1 after treatment with 100 nM HaloBR-6 for indicated amounts of time. Biological triplicates are shown and represented by box and whisker plots. G) Ratio of mClover3-FKBP MFI between tagged cells treated with 20 nM HaloFK-7 or DMSO for 16 hours, in the presence of 5 μM MG132 or vehicle control. Asterisks represent *P* values. H) Ratio of mClover3-FKBP MFI between tagged cells treated with 20 nM HaloFK-7 or DMSO, in the presence of the indicated knockout. 24 and 72 hour treatment was used for WDR26 and RANBP9 cells, respectively. Red points represent unique sgRNAs for knockout. Adjusted *P* values are indicated above bars. * < 0.05, ** < 0.01, *** < 0.001.

We next quantified the degradation of both mClover3-FKBP and BRD4 following WDR26 (Fig. 3E, Fig. S2B) or RANBP9 recruitment (Fig. S2C) by immunoblotting, demonstrating significant degradation of both targets by WDR26 and RANBP9. In comparison, untagged cells or cells with tagged DDB1, an E3 ligase component that was not a hit in the growth screen, did not lead to target degradation in the presence of the appropriate heterobifunctional ligand. Furthermore, we confirmed that BRD4 degradation by WDR26 leads to rapid and near-complete cell depletion while recruitment of the inactive effector, DDB1, to BRD4 has no appreciable effect (Fig. 3F). Similar results are obtained with RANBP9, as quantified by protein levels (Fig. S2D). Based on these results, we conclude that WDR26 and RANBP9 are potent effectors of degradation for multiple targets and thus promising new effectors for targeted protein degradation.

Finally, we explored the mechanisms of degradation for both WDR26 and RANBP9. As expected, the activity of either protein was dependent on the proteasome, as treatment with the proteasome inhibitor MG132 almost completely prevented degradation of mClover3-FKBP (Fig. 3G). To examine the dependency of other CTLH complex subunits for degradation, we generated CRISPR knockouts in WDR26- or RANBP9-tagged cell backgrounds (Table S1). Degradation was carried out for 72 hours in RANBP9-tagged cells, as opposed to 24 hours for WDR26 cells, to increase the signal-to-noise ratio. As a positive control, we showed that the activity of either protein was abrogated (Fig. 3H), and its levels depleted (Fig. S2E), after targeting with CRISPR. To note, RANBP9-targeting sgRNAs downstream of the tag site (Fig. S2F), although functional, did not show significant depletion by flow cytometry most likely due to generation of truncation products (Fig S2E,F). WDR26 activity was also dependent on RANBP9, but RANBP9 activity did not depend on WDR26 (Fig. 3H), reflecting the structural configuration of these proteins in proximity to the catalytic core of the complex^47,48^. Knockout of the catalytic core subunits, MAEA and RMND5A, revealed that while MAEA was necessary for both WDR26- and RANBP9-mediated degradation, RMND5A seemed dispensable for WDR26 but partially required for RANBP9 (Fig. 3G). Although WDR26- and RANBP9-mediated degradation was performed at different time scales, this difference could highlight an additional level of regulation between these subunits. Altogether, these results suggest that the degradation of recruited substrates by WDR26 and RANBP9 occurs through the CTLH complex in a UPS-dependent manner.

## Discussion

Induced protein proximity is poised to transform small molecule therapeutics, yet its full potential will not be realized without scalable methods to profile the effector landscape. Here, we apply our SPOTLITES platform^30^ for pooled endogenous gene tagging with an optimized generic recruitment system to screen for effectors for TPD. Two orthogonal screens reveal multiple potential effectors from different pathways, with members of the CTLH complex, WDR26 and RANBP9, most frequently enriched in this first test of the technology using a focused library. Although RANBP9 was not as potent as WDR26 at inducing degradation, additional optimization of recruitment ligands, linker lengths, binding vectors, and cellular contexts can potentially increase its efficacy. Interestingly, the CTLH complex is frequently upregulated in cancer cells^34^, which could explain why it is most prevalent in our HAP1-based screens. This cancer-specific upregulation, in addition to recent evidence implicating multiple CTLH subunits, particularly WDR26, in tumorigenesis^49^, makes WDR26 and RANBP9 excellent candidates for new TPD cancer therapies that could have both increased tissue specificity and reduced susceptibility to resistance mutations as compared to existing degraders^18^. Future work will focus on the development of ligands for effectors validated via this approach to expand the toolbox of E3 ligase ligands available for TPD.

In this work, we focused on screening select members of the proteostasis network for effectors of TPD using two target proteins within a single cellular context. The scalability of our approach provided a platform technology which will enable us to easily expand to other cell lines and targets, increasing the toolbox of effectors for TPD and characterizing their target specificities and activities across cell lines. Future tagging libraries can easily expand the potential effector space beyond members of the proteostasis network to explore other forms of post-translational protein modifications within a diverse range of cellular phenotypes in a systematic, biology forward fashion, as opposed to the chemistry forward approaches employed to date in the TPD field. The protein effectors that will be discovered through these methods could be harnessed to study protein function, dramatically improve the potency of current TPD efforts, and expand induced proximity approaches to new therapeutic modalities. Crucially, this can all now be performed rapidly with endogenous proteins, in a scalable fashion, with minimal upfront bespoke chemical probe development, enabling the selection of the optimal target-effector pair.

## Acknowledgements

We would like to thank the Shalem and Burslem labs for extensive discussions related to this manuscript. We thank the Prof. Craig Crews lab for sharing reagents (PhosTAC7 and GFP-FKBP HEK293s). This work was supported by the following grants: DP2GM137416 from NIH/NIGMS, SAP#4100083086 from PA DoH and R03NS111447-01 from NINDS awarded to O.S. R35GM142505 from NIGMS awarded to G.M.B. F32CA239499 from NCI and K99AG075256 from NIA awarded to Y.V.S.

## Declaration of interests

Y.V.S., G.M.B., and O.S. have filed several patent applications related to this manuscript through the Children’s Hospital of Philadelphia and the Perelman School of Medicine at the University of Pennsylvania. G.M.B serves on the scientific advisory board of Plexium Inc. and has received consulting payments from Chinook Tx., Repare Tx. and Intima Biosciences Inc. None of these entities have input into or prior knowledge of the studies reported here.

## Methods

### Antibodies

The following antibodies were used for immunoblotting: BRD4 (NBP2-76393, Novus Biologicals; 1:1000), GFP (sc-9996, Santa Cruz Biotechnology; 1:1666), alpha-tubulin (NB100-690, Novus Biologicals; 1:5000), IRDye 680LT Goat anti-Rabbit (LI-COR 926-68021, 1:10,000), IRDye 800CW Goat anti-Mouse (LI-COR 926-32210, 1:10,000).

### Immunoblotting

Cultured cells were pelleted, washed with PBS, and resuspended in RIPA lysis buffer (Cell Signaling, 9806) with 1x protease inhibitor cocktail (MilliporeSigma, P8340). Samples were normalized by bicinchoninic acid (BCA) assay (Cell Signaling, 7780), and loaded on a precast SDS-PAGE gel (Bio-Rad, 4561086). Western blotting followed using standard protocols. Imaging of blots was performed on a LI-COR Odyssey instrument.

### Cell culture

Cell lines, including HAP1 (Horizon Discovery) and HEK293 (ATCC CRL-1573) cells were handled according to manufacturer’s instructions. HEK293 cells expressing EGFP-FKBP^29^ were a gift from the Craig Crews lab. Clonal HAP1 haploid lines were generated by single cell sorting and genome editing was performed within 1-2 weeks of expansion to ensure most cells were haploid during editing. HAP1 cells were cultured in IMDM (Gibco, 12440053) + 10% fetal bovine serum (Thermo Fisher Scientific, FB12999102) + 1% antibiotic-antimycotic (Gibco, 15240062). HEK293 cells were cultured in DMEM (Gibco, 11995065) + 10% fetal bovine serum (Thermo Fisher Scientific, FB12999102) + 1% antibiotic-antimycotic (Gibco, 15240062). All cells were dissociated with TrypLE Express (Thermo Fisher Scientific, 12605010).

### Lentiviral transduction

Transduction was performed for generating cells expressing the mClover3-FKBP target or for CRISPR KOs (Table S1). mClover3-FKBP was expressed from an EF1a-driven transfer vector (Addgene TBA), and sgRNAs and Cas9 were expressed from lentiCRISPR v2 (Addgene 52961). Virus was generated in HEK293T cells by transfection with the aforementioned transfer vectors and pMDLg (Addgene 12251), pRSV-REV (Addgene 12253), and pMD2g (Addgene 12259). mClover3-FKBP-containing virus was transduced at low efficiency (∼10%) as measured by flow cytometry, while lentiCRISPR v2 transduction efficiency was ∼50% as measured by cell counting after puromycin selection. Infection was performed in the presence of polybrene (Millipore, TR-1003-G), a day after which the cells were replenished with fresh media. Enrichment was performed 48 hours after infection by fluorescence-activated cell sorting (FACS) or puromycin selection, as appropriate.

### Ligand treatments and competition assays

HaloTag-TMR (Promega, G8251) was used to fluorescently label tagged cells. Cells were treated with the ligand at 20 nM final concentration for 15 minutes at 37°C with 5% CO2. Cells were then washed 3x with fresh media and incubated for an additional 30-60 minutes in media. Labeled cells were analyzed either by microscopy or flow cytometry.

dTAG^V^-1 (Tocris Bioscience, 6914) was used to degrade FKBP^F36V^-tagged proteins at the indicated concentrations and timescales (typically ∼24 hours).

Recruitment of tagged proteins was performed with either HaloFK-2,-5,-7, or HaloBR-6 with the indicated concentrations and timescales. Compound synthesis is described in the Supplementary Text. An aliquot of HaloFK-7 (i.e. PhosTAC7^9^) was provided as a gift from the Craig Crews lab.

For HaloFK-2 competition experiments, cells were pretreated with HaloFK-2 for ∼16 hours before co-treatment with the indicated concentrations of HaloTag-TMR for 15 minutes or dTAG^V^-1 for 10 hours.

MG132 (MilliporeSigma, M7449) was used to determine the role of the proteasome in TPD. Cells were pre-treated with 5 μM MG132 or vehicle control for 4 hours before co-treatment with 20 nM HaloFK-7 for 16 hours.

### Confocal imaging

Cells were grown on coverslips and directly fixed in 4% formaldehyde (Electron Microscopy Sciences) in PBS (Thermo Fisher Scientific). Fixed cells were washed in PBS and coverslips were mounted on microscopy slides in ProLong Glass Antifade Mountant with NucBlue Stain mounting medium (Thermo Fisher Scientific, P36985). Images were acquired on a Leica TCS SP8 confocal microscope. Z-stacks (0.6 μm slices) spanning the entire volume of the cells were recorded with oil-immersion 63x Plan-Apochromat lenses, 1.4 NA.

### Flow cytometry and cell sorting

Cultured cells were trypsinized, resuspended in media to ∼1×10^6^ cells/ml, and filtered through a cell strainer. Cellular fluorescence was measured on a BD FACSAria Fusion (BD Biosciences). All cells were analyzed or sorted using an 85 μm nozzle. HaloTag-TMR fluorescence was detected by the 561 nm laser and the 582/15 filter. GFP fluorescence was detected by the 488 nm laser and filters 502LP and 530/30. “Autofluorescence” was detected by the 405 nm laser and the 450/50 filter. Data were analyzed using the R package flowCore (v2.2.0). For quantification of degradation, the mClover3 fluorescence baseline was determined using non-fluorescent parental cells when available, or with dTAG^V^-1-treated samples otherwise.

Polyclonal sorting was performed according to cytometer instructions into 15 ml conical tubes at 4°C partially filled with media. To culture sorted cells, tubes were centrifuged at 300 g for 3 min and the cells were resuspended in fresh media and then transferred to a tissue culture plate. Single cell sorting was performed directly into 96-well tissue culture plates containing 50 μl media.

### DNA content analysis

Cells were cultured in media containing 5 μg/ml Hoechst 33342 Solution (Thermo Fisher Scientific, 62249) at 37°C for 30 - 60 minutes. Cells were then immediately lifted and resuspended in PBS on ice. DNA content was measured using the flow cytometer with the 405 nm laser and the 450/50 filter.

### Viability assays

For assessing cell viability by counting, thoroughly resuspended cells were analyzed on the Countess II (ThermoFisher) according to manufacturer’s instructions. The MTS or CellTiter 96 AQueous One Solution Cell Proliferation Assay (Promega, G3580) was performed in transparent 96-well tissue culture plates (Corning, 3596). The reagent was warmed to room temperature and 20 μl was added to cells in 100 μl media, which were then incubated in a tissue culture incubator for 1 to 4 hours. Absorbance at 490 nm was then measured on a SpectraMax i3 plate reader.

### Pooled tagging: library design

Genes of interest were selected using the following Gene Ontology^50^ annotations: ubiquitin ligase complex, protein ubiquitination, autophagy, protein folding, and misfolded protein binding. From the list of genes, the intron-targeting sgRNA library was generated as described previously^30^, or via the www.pooledtagging.org online database.

### Pooled tagging: plasmid library generation

In generating pooled tag cell libraries, the protocol for “plasmid library generation” and “viral library generation and transduction” (next section) follows typical CRISPR library generation methods^39^. Nonetheless, we present all steps in detail below.

1. An oligo pool was synthesized by Twist Bioscience or as oPools Oligo Pools (IDT) with the following design logic: 5’ - fw. primer (N20) - BsmBI binding (CGTCTC) - backbone overhang (ACACCG) - spacer seq. (N20) - backbone overhang (GTTTT) - BsmBI binding (GAGACG) - rev. primer (N20) - 3’
2. Oligos were PCR amplified and purified. 25 μl reactions were prepared for each sublibrary with the KAPA HiFi PCR Kit (Roche, KK2502) containing 0.5 μl of the oligo pool resuspended to 20 ng/μl, 5 μl 5x KAPA HF buffer, 0.75 μl 10 mM dNTP mix, 0.5 μl polymerase, 0.75 μl appropriate 10 μM primer mix. PCR cycling parameters: 1) 3 min. at 95°C, 2) 20 sec. at 98°C, 3) 15 sec. at 65°C, 4) 15 sec. at 72°C, 5) repeat steps 2-4, 10x, 6) 1 min. at 72°C. PCR products were gel purified with the Monarch Gel Extraction Kit (NEB, T1020).
3. Amplified oligos were ligated into an sgRNA-expressing backbone by Golden Gate cloning with BsmBI (NEB, R0739) and T7 ligase (NEB, M0318), using 5 ng of PCR product and 50 ng of the CROPseq backbone (Addgene, 86708). PCR cycling parameters: 1) 5 min. at 42°C, 2) 5 min. at 16°C, 3) repeat steps 1-2, 50x, 4) 10 min. at 65°C.
4. Plasmids were transformed and purified. Golden Gate products were first concentrated by IPA precipitation. 1 volume sample was mixed with 1 volume isopropanol in 50 mM NaCl and incubated at room temperature for 15 minutes. Samples were then centrifuged at 15 k x g for 15 minutes at room temperature and washed 2x with cold 70% EtOH. After drying the DNA pellets, samples were resuspended in 5 μl H_2_O. Electroporation was performed according to manufacturer’s instructions into Endura ElectroCompetent Cells (Lucigen, 60242-2). Dilution of transformed cells and estimation of the number of colony forming units allowed for calculation of plasmid library coverage of around 200x. Bacteria was cultured overnight and plasmids were purified with a maxiprep kit according to manufacturer’s instructions (Thermo Fisher Scientific, A31217).

### Pooled tagging: viral library generation and transduction

1. Lentivirus was produced from HEK293T cells seeded into a 15 cm tissue culture dish at 10 M cells / dish. Transfection was performed with PEI (Polysciences, 24765) in Opti-MEM (Thermo Fisher Scientific, 31985062). 136.35 μl PEI (3 μl PEI : 1 μg DNA) was incubated in 1.25 ml Opti-MEM, and a separate solution of 1.25 ml Opti-MEM was mixed containing: 1) 20 μg plasmid pool, 2) 13.25 μg pMDLg, 3) 7.2 μg pMD2g, 4) 5 μg pRSV-REV. Both solutions were combined, incubated for 15 minutes, and 2.5 ml added dropwise to cells containing 15 ml of antibiotic-free media. Media was replenished the following day, and viral aliquots were collected 48 hours post-transfection, filtered through 0.45 μm cellulose acetate filters (VWR, 76479-040) or 0.45 μm PES filters (Thermo Fisher Scientific, 50-607-518), and frozen at -80°C.
2. Virus was titered and cells were transduced. Titering was performed by puromycin selection and cell counting. HAP1 cells expressing mClover3-FKBP were infected with the lentiviral library at ∼30% transduction efficiency and in amounts to achieve 1000 transduced cells per sgRNA. Virus was administered by “spinfection”, where cells and virus were mixed with polybrene (Millipore, TR-1003-G) in antibiotic-free media at 2 M cells per well of a 12-well plate. Plates were spun at 32°C for 1 hour at 1000 x g and placed in a tissue culture incubator overnight. The following day, cells were expanded into 15 cm dishes in the presence of 1 μg/ml puromycin (Gibco, A1113803), and selection took place over 5 days.

### Pooled tagging: tagged cell library generation

1. Transduced cells were transfected with reagents necessary for homology-independent tagging. Based on an assumed average tagging efficiency of 0.05% in HAP1 cells, we transfected enough cells for 10x coverage for each tagging library. Cells were plated at ∼13 M cells per 15 cm dish in 15 ml of antibiotic-free media. Transfection was performed with TurboFectin 8.0 (OriGene, TF81001). 85 μl reagent (3 μl reagent : 1 μg DNA) was incubated in 1.25 ml Opti-MEM, and a separate solution of 1.25 ml Opti-MEM was mixed containing: 1) 8.6 μg tag donor pMC_*X*_BSDminus_*Y* (where *X* refers to the fusion domain and *Y* refers to the sequence phase of the library), 2) 6.1 μg sgDonor_lentiGuide, and 3) 13.5 μg lentiCRISPR. The molar ratio of tag donor to any other plasmid was kept 5 : 1. The tag donor was also introduced as a minicircle generated with the MC-Easy Minicircle DNA Production Kit according to manufacturer’s instructions (System Biosciences, MN920A-1). Both solutions were combined, incubated for 15 minutes, and added to cells dropwise. Media was replenished after 5 - 6 hours of incubation.
2. Cells with successful genomic integration of the tag donor were selected using the blasticidin resistance cassette^40^. 48 hours after transfection, media with 15 μg/ml blasticidin (Gibco A1113903) was added to cells. Cells were incubated for 3 days until all untransfected cells depleted. Cells were subsequently expanded for 3 days in the presence of 5 μg/ml blasticidin. After expansion, cells were split in the presence of 15 μg/ml blasticidin and another round of selection occurred, this time between cells with and without genomic integration of the donor. Again, cells were expanded in the presence of 5 μg/ml blasticidin until they were ready for sorting.
3. Cells with expressed endogenous fusion proteins were purified by fluorescence-activated cell sorting (FACS) as described in the “Flow cytometry and cell sorting” section. Prior to sorting, HaloTag was labeled with HaloTag-TMR (Promega, G8251) as described in the “Drug treatments and competition assays” section. 3 - 5 M cells per library were sorted, based on library size. A second round of FACS was performed for each library to ensure the percentage of HaloTag-expressing cells was at least 95%.

### Pooled tagging: tagged library amplicon sequencing

Tag library composition was determined by bulk sequencing of sgRNA amplicons. Genomic DNA was extracted with the QIAamp DNA Mini Kit (QIAGEN, 51304) according to the manufacturer’s instructions. Extracted gDNA was PCR amplified (Takara, RR001A) using a 30-cycle protocol. Each reaction had 5 μg of gDNA template. PCR products were purified by gel extraction (NEB, T1020) and sequenced on a NextSeq 500 (Illumina) with a 5% spike-in of PhiX v3 (Illumina, FC-110-3001).

### Fluorescence screen

Tagged cells were treated with 20 nM HaloFK-7 in duplicate or with HaloFK-2 for 22 hours. Cells were trypsinized and collected for FACS on ice. Sorting gates were drawn as described in Figure S1E, and cells were sorted at 4°C to ∼4M cells per sorting gate. Cell counts were determined by the Countess II (ThermoFisher). Subsequently, cells were pelleted and stored at - 20°C until genomic DNA extraction was performed as described in “*Pooled tagging: tagged library amplicon sequencing*”.

### Growth screen

Tagged cells were treated with 100 nM HaloBR-6 or DMSO for 72 hours in triplicate. Each sample was maintained at a 15 M cell bottleneck to ensure that >90% of sgRNAs had > 50x coverage, with an average coverage of ∼1800x. Subsequently, cells were pelleted and stored at -20°C until genomic DNA extraction was performed as described in “*Pooled tagging: tagged library amplicon sequencing*”.

### Analysis of fluorescence and growth screens

sgRNA sequences were extracted from sequencing reads, aligned to a reference dataset of all possible sgRNAs, and demultiplexed using the R package *CB2* (v1.3.4). sgRNAs were filtered to remove the minor mode of count distributions as shown in Figure S1B. Subsequent count normalization, sgRNA fold changes, and FDRs were calculated using the R package *edgeR* (v3.32.1).

## Notes

### Competing Interest Statement

Y.V.S., G.M.B., and O.S. have filed several patent applications related to this manuscript through the Childrens Hospital of Philadelphia and the Perelman School of Medicine at the University of Pennsylvania. G.M.B serves on the scientific advisory board of Plexium Inc. and has received consulting payments from Chinook Tx., Repare Tx. and Intima Biosciences Inc. None of these entities have input into or prior knowledge of the studies reported here.

