## Supplementary Figures for "Pooled endogenous protein tagging and recruitment for scalable discovery of effectors for induced proximity therapeutics"

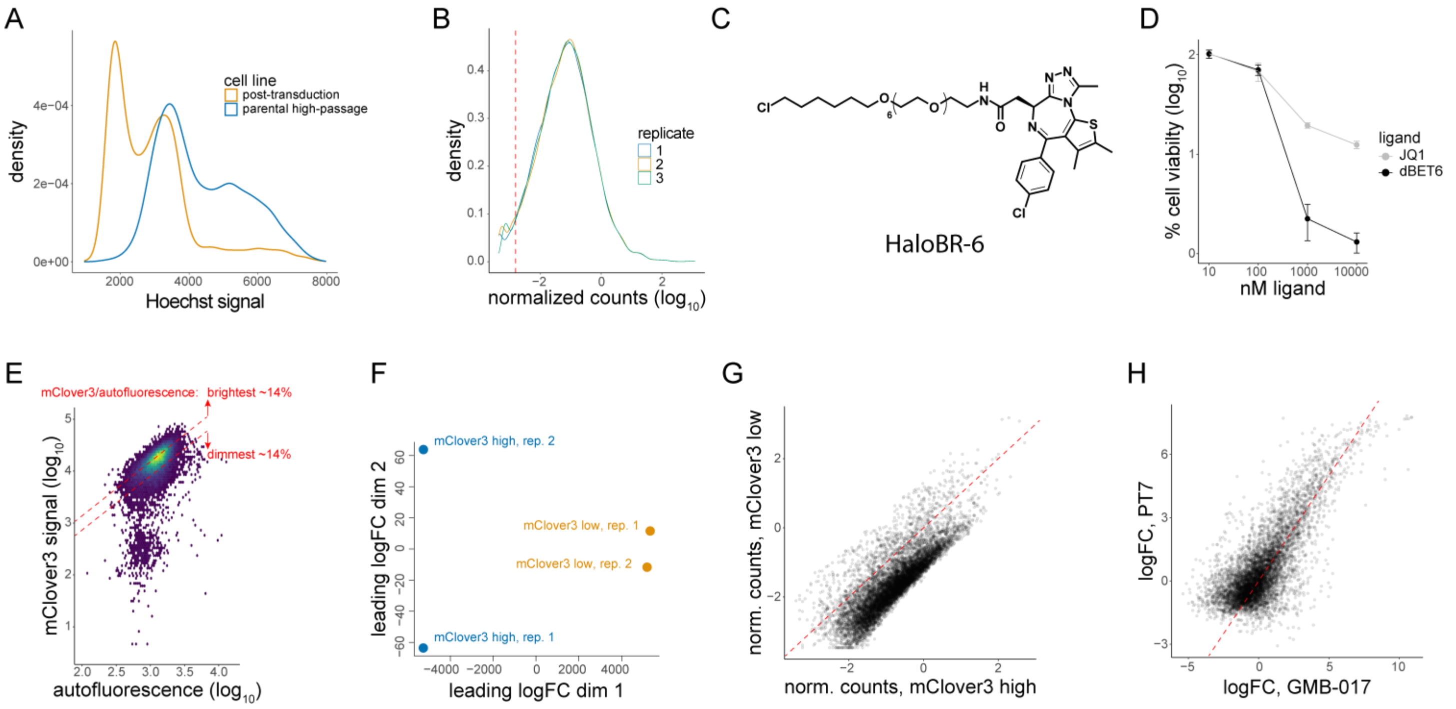


**Figure S1. Constructing and utilizing a pooled tag library for screening effectors for TPD, related to Figure 2.** A) DNA content analysis by flow cytometry of HAP1 cells after transduction of sgRNAs for tagging but before transfection of tagging reagents. B) Density plot of normalized counts of sgRNA amplicons from tagged libraries in three replicates. The red dashed line represents the threshold for detected sgRNAs. C) Chemical structure of HaloBR-6. D) MTS cell viability assay in HAP1 cells treated with a BRD4 inhibitor (JQ1) or degrader (dBET6) for 72 hours. E) Flow cytometry scatter plot of pooled tag library. Red dashed lines represent gates for mClover3-FKBP high- or low-expressing cells. F) MDS plot for samples from a pooled tag library treated with HaloFK-7 for 24 hours and sorted on mClover3 fluorescence. G) Scatter plot of sgRNA counts from mClover3-FKBP high- or low-expressing populations in a pooled tag library treated with HaloFK-7 for 24 hours. H) Scatter plot of sgRNA mClover3 low/high log fold changes between cell lines treated with HaloFK-7 or HaloFK-2 for 24 hours.

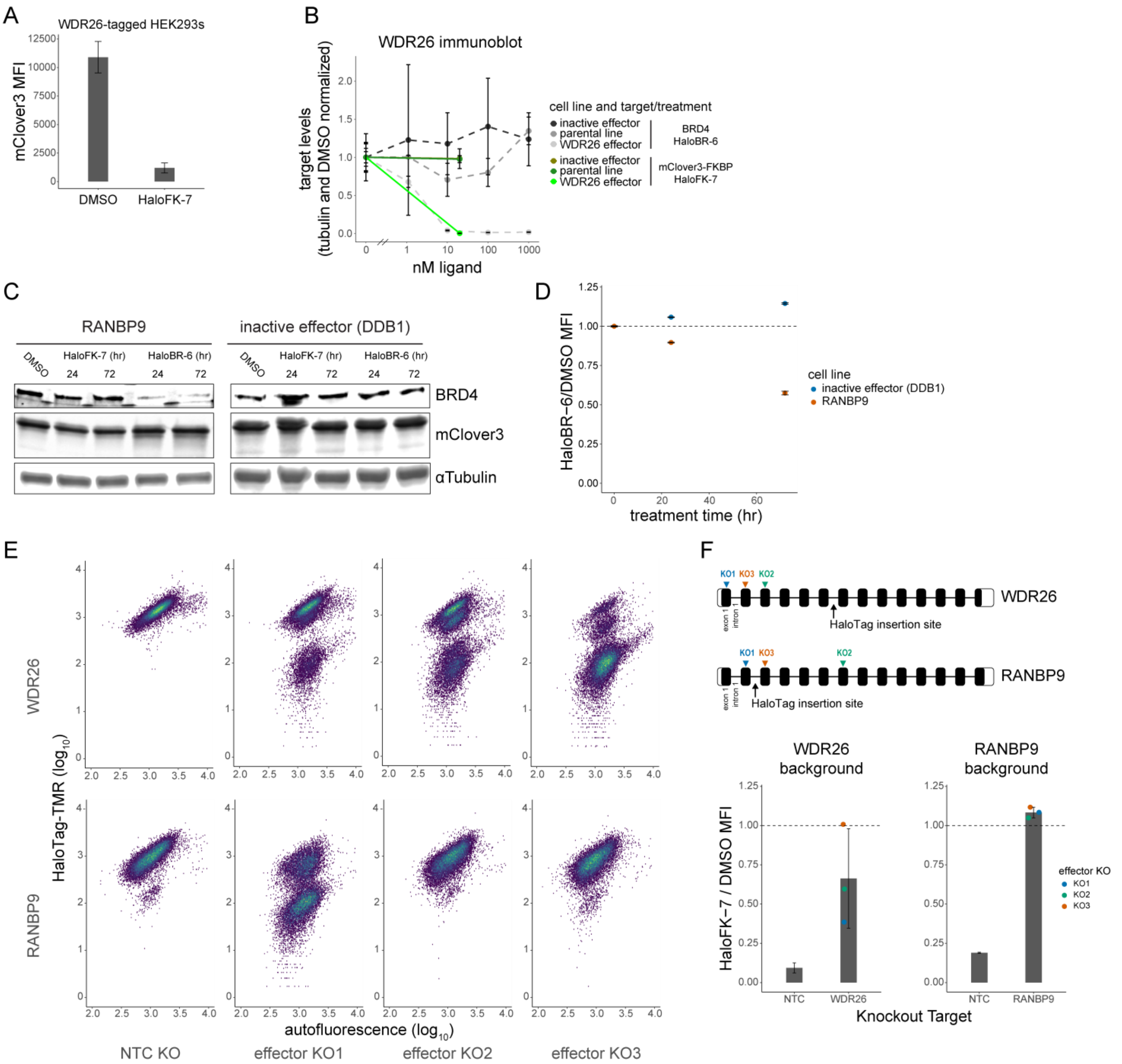


**Figure S2. Characterization of new effectors for TPD, related to Figure 3.** A) Mean fluorescence intensity (MFI) of EGFP-FKBP$^{F36V}$ in HEK293 cells with endogenously tagged WDR26 treated with HaloFK-7 for 24 hours, as measured by flow cytometry. B) Quantification of immunoblot from Figure 3E. C) Immunoblot representing 24 or 72 hour degradation of BRD4 or mClover3-FKBP with 100 nM HaloBR-6 or 20 nM HaloBR-7, respectively, in cells with tagged WDR26 or the DDB1 inactive effector. D) Total protein levels, representing cell growth, from RANBP9- or DDB1-tagged cells treated with 100 nM HaloBR-6 for the indicated times. E) Flow cytometry of HaloTag-TMR-labeled cells with tagged WDR26 or RANBP9 and treated with non-targeting control (NTC) sgRNAs, or sgRNAs against the tagged effector. F) Depiction of sgRNA target sites and activities. Related to Figures 3H and S2E.
