## Supplementary Text for "Pooled endogenous protein tagging and recruitment for scalable discovery of effectors for induced proximity therapeutics"

### Synthesis of HaloFK-2,-5,-7

HaloFK-2,-5 and -7 were synthesized according to literature procedures and matched reported analytical data (*ACS Chem. Biol.* 2021, 16, 12, 2808–2815).

### Synthesis of HaloBR-6

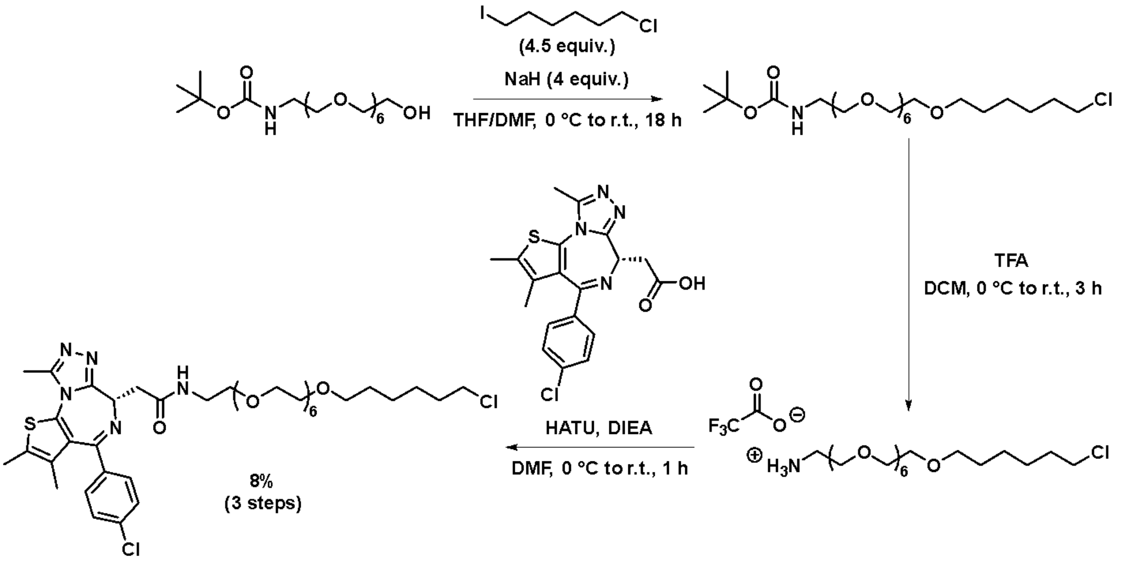


Step 1

To a solution of *N*-Boc-PEG$_7$-OH (1 equiv., 100 mg, 0.24 mmol) in THF (1.3 mL) and DMF (0.7 mL) at 0 °C added portionwise NaH (4 equiv., 60% dispersion in mineral oil, 37 mg, 0.91 mmol). After stirring at 0 °C for 30 min (or until no bubbling can be observed), 6-chloro-1-iodohexane (4.5 equiv., 0.16 mL, 1.02 mmol) was added to the mixture at 0 °C. The reaction mixture was stirred at 0 °C and let to come back to room temperature overnight. The reaction was then quenched at 0 °C with 2 mL of MeOH and the volatiles were evaporated. The residue was chromatographed on silica gel (DCM to 2.5% MeOH in DCM) to afford the product as a clear oil (125 mg).

Step 2

To the product obtained above (125 mg) dissolved in dichloromethane (3 mL) at 0 °C was added dropwise trifluoroacetic acid (0.8 mL). The mixture was then stirred at room temperature for 4 hours. Solvents were removed *in vacuo.* Then 3 mL of toluene were added to the residue to azeotropically evaporate residual TFA. 3 mL of $Et_2O$ were finally added and evaporated to afford the crude product (80 mg) that was used in the next step without further purification.

Step 3

JQ1-COOH (1.2 eq., 69 mg, 0.17 mmol) was dissolved in DMF (0.67 mL) followed by the crude obtained above (1 eq., 80 mg, 0.14 mmol) and DIEA (5.0 eq., 0.13 mL, 0.72 mmol). HATU (2 eq., 109 mg, 0.29 mmol) was added and the reaction mixture was stirred for 1 h. Volatiles were concentrated and the crude was purified first by preparative reversed-phase HPLC chromatography (Agilent 1260 Infinity II HPLC system) on a Agilent Prep-C18 column (250 mm × 21.2 mm × 10 µm) using a linear gradient (5% to 95% in 24 min, flow-rate of 20 $mL.min^{-1}$) of solvent B (0.1% TFA in ACN, v/v) in solvent A (0.1% TFA in $H_2O$, v/v). UV detection was set at 254 and 280 nm. The volatiles were then evaporated in a SpeedVac Concentrator (Fisher Scientific) but analyses showed the presence of another product. This impurity was associated with the hydrated product probably due to the low pH of the solvents used during the purification. All fractions were then gathered and purified by silica gel chromatography (0% to 9% MeOH in DCM) to give the corresponding product as a light-yellow oil (17 mg, 8% over three steps).

**$^1$H NMR** (400 MHz, CDCl3) δ 7.42 – 7.37 (m, 2H), 7.34 – 7.29 (m, 2H), 6.89 (t, J = 5.5 Hz, 1H), 4.64 (t, J = 7.0 Hz, 1H), 3.72 – 3.32 (m, 34H), 2.65 (s, 3H), 2.39 (d, J = 0.9 Hz, 3H), 2.04 (bs, 2H), 1.81 – 1.71 (m, 2H), 1.66 (d, J = 0.9 Hz, 3H), 1.63 – 1.53 (m, 2H), 1.50 – 1.29 (m, 4H). **$^{13}$C NMR** (101 MHz, CDCl3) δ 170.6, 163.8, 155.7, 149.8, 136.7, 132.2, 130.9, 130.7, 130.5, 129.9, 128.7, 71.2, 70.6, 70.6, 70.6, 70.4, 70.1, 69.8, 54.4, 45.1, 39.5, 39.1, 32.6, 29.5, 26.7, 25.4, 14.4, 13.1, 11.8. **LC/MS:** Calculated for $C_{39}H_{58}Cl_2N_5O_8S$ $[M+H]^+$: 826.3 ;Found: 825.4. Calculated for $C_{39}H_{59}Cl_2N_5O_8S$ $[M+2H]^{2+}$: 413.7; Found: 413.3.
